# Proximity Labelling Reveals Spatial Organisation of Mitochondrial Protein Import Complexes

**DOI:** 10.64898/2026.07.20.739561

**Authors:** Xu Wang, Claudia Cavarischia Rega, Boris Maček, Ralf-Peter Jansen

## Abstract

Mitochondrial protein import and local translation at the mitochondrial outer membrane (MOM) require coordinated interactions between protein translocases and RNA-associated factors, yet the molecular organisation of these interactions remains largely unresolved. Here, we used APEX2-mediated proximity labelling to define the proximal proteomes associated with the cytosolic face of the TOM and SAM complexes by fusing APEX2 to TOMM22 and MTX2, respectively. Quantitative mass spectrometry identified known TOM/SAM-associated proteins together with multiple RNA-binding proteins (RBPs), supporting the emerging role of the MOM in localised translation and other RNA-related processes. Among identified RBPs, the exonuclease EXD2 was consistently enriched in APEX2-TOMM22 and MTX2-APEX2 datasets and remained associated upon puromycin treatment, indicating a translation-independent interaction. Together, our findings provide insights into the molecular organisation of mitochondrial import sites and identify EXD2 as a TOM/SAM-associated factor at the MOM.

## Introduction

Mitochondria are essential double-membraned organelles whose functions rely on the nuclear genome to encode approximately 99% of their proteome^1, 2^. Nuclear-encoded proteins are synthesized on cytosolic ribosomes and must be accurately targeted and imported into the organelle through specialized protein machineries ^3, 4^. The translocase of the outer membrane (TOM) complex serves as the general entry gate for nearly all nuclear-encoded mitochondrial proteins. This complex consists of receptor proteins—primarily Tom20 (TOMM20 in mammals), Tom22/TOMM22, and Tom70/TOMM70—which recognize diverse targeting signals, and the pore-forming protein Tom40/TOMM40, which allows preproteins to traverse the outer membrane into the intermembrane space ^3, 5^.

While the TOM complex facilitates translocation, the stable insertion of β-barrel proteins, including the metabolite porins or Tom40 itself, into the outer membrane is the primary task of the sorting and assembly machinery (SAM) complex, also known as the TOB complex ^3, 6–8^. The SAM complex is an evolutionarily ancient machinery composed of the channel-forming subunit Sam50 along with peripheral capping proteins Sam35 (MTX2 in mammals) and Sam37/MTX1 ^9^.

It has been established that the TOM and SAM complexes are physically and functionally connected within a dynamic protein network rather than operating as independent units ^3, 6^. The two machineries can associate to form a TOM-SAM supercomplex, which facilitates the efficient transfer of β-barrel precursors directly from the TOM import channel to the SAM insertion sites ^10^. The subunit Sam37/MTX1 plays a critical role in this coupling, acting as a bridge that promotes precursor transfer and stabilizes the formation of the nascent β-barrel ^11^. However, based on the estimated number per yeast cell of TOM complex subunits (approx. 20,000) and SAM complex subunits (approx. 1,500), it is likely that a large portion of TOM complexes must be present in a SAM-free state ^12, 13^. SAM-associated and SAM-free complexes might thus be differentially compartmentalized in the mitochondrial outer membrane (MOM). Furthermore, TOM and SAM complexes might associate with different cytoplasmic partners such as chaperones or RNA-associated factors, including ribosomes ^14, 15^. This might be of special importance for translation-coupled protein import into mitochondria ^16^, which has been shown to occur for large multi-domain mitochondrial proteins ^17, 18^. Furthermore, several RNA-binding proteins (RBPs) that associate with mitochondrial protein-encoding mRNAs are found at or close to the mitochondrial surface, including yeast Puf3 ^19^, or human AKAP1 ^18, 20^ and SYNJ2a ^21^. For some of these RBPs, an association with the TOM complex has been suggested ^22, 23^. If such an association with the SAM complex occurs, has yet to be shown.

To probe the molecular environment of TOM and SAM complexes in living cells, we performed proximity labelling ^24, 25^ using the central receptor of the TOM complex (TOMM22) and the scaffolding protein MTX2 of the SAM complex as baits. As we were particularly interested in cytoplasmic interactions, we fused the cytoplasmic face of each protein with the APEX2 proximity labelling enzyme ^26^. Here, we show that overlapping yet distinct sets of cytoplasmic RBPs are enriched in the proximity of both TOM and SAM complexes in a translation-independent manner, including the exonuclease EXD2 that has been previously implicated in resolving transcriptional stalling in the nucleus.

## Material and Methods

### PCR and Cloning

PCR reactions were performed with Herculase II Fusion DNA Polymerase (Agilent, #600675) according to the manufacturer’s protocol. For cloning, Gibson Assembly was performed for the sequence- and orientation-specific ligation between the insert and the plasmid backbone. For a 20 μL reaction, 10 μL 2× NEBuilder® HiFi DNA Assembly Master Mix (NEB, #E2621L) was added to 10 μL of diluted DNA fragments (100 ng plasmid backbone with 3-fold molar excess of insert). The reaction mixture was incubated at 50 °C for 1 h in a PCR machine before transformation into Top10 *E. coli* through heat shock. All plasmids generated in this study are listed in **Supporting Table 1**.

### Cell culture

Wild-type HeLa-EM2-11ht cells were cultured in Dulbecco’s Modified Eagle’s Medium (Sigma-Aldrich, #D6046-500ML) supplemented with 10% fetal bovine serum, 200 μg/mL G418, and 1× penicillin / streptomycin at 37 °C with 5% CO_2_.

### Generation of cell lines stably expressing APEX2-fusion constructs

For stable cell line generation, 150,000 HeLa-EM2-11ht cells were seeded per well in a 6-well plate 24 h before transfection. At approximately 60% confluency, cells were co-transfected with 2 μg pSF3 plasmid containing the corresponding APEX2 fusion construct and 2 μg pPGKFLPobpA plasmid (RJP2042) using FuGENE® HD Transfection Reagent (Promega, #E2311) with a 3:1 ratio (transfection reagent to plasmid DNA, v/w) in 500 μL Opti-MEM. Culture medium was replaced 24 h after transfection and 50 μg/mL ganciclovir was applied 72 h after transfection and was kept for 14 days with medium replacement every 48 h. For monoclonal cell line generation, cells were serial diluted in a 96-well plate, and wells containing only one clone were selected. After expansion, each clone was screened by Western blotting for expression of the desired APEX2 fusion construct.

### RNA Interference

In the design of the APEX2-TOMM22 construct, we introduced 4 silent mutations in exon 3 of TOMM22 (within positions 370-382, NCBI RefSeq: NM_020243.5) to avoid targeting and degradation by the siRNA. For the knockdown of endogenous TOMM22, 150,000 cells were seeded per well in a 6-well plate and supplemented with 300 ng/μL doxycycline 24 h before siRNA transfection to allow expression of the APEX2-fused construct. 10 nM Silencer™ TOMM22 siRNA (siRNA ID: 133043, Invitrogen) and 5 μL Lipofectamine 3000 (Invitrogen, #L3000008) were separately diluted in 100 μL Opti-MEM, incubated for 5 min at RT, and pooled together. After mixing, the transfection mixture was incubated at RT for another 15 min before adding to the 6-well plate. For the negative control, 10 nM Silencer™ Select Negative Control No. 1 siRNA (#4390843, Invitrogen) was used. After transfection, cell culture medium and doxycycline were replaced every 24 h, and cells were harvested 72 h post-transfection for Western blot analysis.

### Isolation of crude mitochondria

Cells were collected from a 15 cm dish and washed with PBS. Each pellet was resuspended in 1 mL ice-cold mitochondrial isolation buffer (MIB, 70 mM sucrose, 210 mM mannitol, 5 mM HEPES, 1 mM EGTA) and lysed by sequentially passing through 20G, 25G, and 27G needles. The lysate was incubated on ice for 10 min before spinning down at 900 × g for 5 min at 4 °C. The supernatant was collected and centrifuged again at 9,000 × g for 20 min at 4 °C to pellet crude mitochondria. After removing the supernatant, the pellet was washed twice with 1 mL MIB and resuspended in 200 μL MIB. Protein concentration was determined using a Nanodrop instrument.

### Protein analysis

For all Western blots, 30-50 μg of protein was loaded in each well and separated on 12% SDS-PAGE gels. Subsequently, proteins were blotted onto either nitrocellulose or PVDF membranes. Ponceau S staining was performed for most nitrocellulose membranes. Blots were blocked in 5% milk (w/v) in TBST (Tris-buffered saline, 0.1% Tween-20) at RT for 1 h. Primary antibody (**Supporting Table 2**) incubation was performed in 5% milk in TBST for either 1 h at RT or overnight at 4 °C. After 3 × 10 min washes in TBST, blots were incubated with secondary antibody in 5% milk in TBST for 1 h at RT. Blots were washed 3× 10 min in TBST before development with the Pierce^TM^ ECL Western Blotting substrate (Thermo Fisher), and images were acquired on the ChemiDoc Imaging System (Bio-Rad). For probing with streptavidin-HRP, only nitrocellulose membranes were used and membranes were blocked in 3% BSA (w/v) in TBST for 1 h at RT and incubated with 0.5 μg/mL streptavidin-HRP in 3% BSA for 1 h at RT.

For the analysis of TOMM22 integration in the TOM complex, blue-native (BN)-PAGE was used ^27^. Crude mitochondria were isolated, and the MOM was solubilised with digitonin (6:1 digitonin-to-protein ratio, w/w) at 4 °C for 1 h on a rotating wheel. The insoluble mitoplast fraction was removed by centrifugation at 13,000xg for 30 min at 4 °C and the solubilised MOM fraction was collected and loaded on a 6-14% gradient gel. After blotting onto a PVDF membrane, it was handled in the same way as a Western blot.

### Co-immunoprecipitation

Mitochondria were isolated in MIB supplemented with 1× protease inhibitor cocktail (Roche, #11873580001). 1 mg of isolated mitochondria was used per sample and was solubilised with digitonin as described above. The supernatant containing solubilised MOM was transferred to another microcentrifuge tube with 100 μL anti-V5 magnetic agarose beads (Proteintech, #v5tma) equilibrated twice with MIB. Co-immunoprecipitation was performed at room temperature for 1 h on a rotating wheel. After three washes with MIB, proteins were eluted by boiling at 95 °C for 15 min in 2× SDS-PAGE sample buffer.

### Immunofluorescence

Cells were seeded on glass coverslips in 12-well plates and induced with doxycycline 24 h before the experiment. Prior to immunostaining, cells were fixed with 4% paraformaldehyde in PBS for 15 min followed by three washes with PBS. Cells were permeabilised in 0.5% Triton X-100 in PBS for 15 min. After three washes with PBS, cells were blocked in 3% BSA overnight at 4°C. Primary antibody incubation (1:200 rabbit anti-TOMM20 and 1:1,000 mouse anti-V5 for APEX2-TOMM22, MTX2-APEX2, and APEX2-MAVS; 1:1,000 diluted mouse anti-V5 for APEX2-NES) was performed in 3% BSA for 1 hour at 37 °C. Following 3 × 5 min washes with PBS, cells were incubated with secondary antibodies (Alexa Fluor goat anti-mouse-488, 1:200; Alexa Fluor goat anti-rabbit-350, 1:100) in 3% BSA at 4°C overnight. Coverslips were mounted with Vectashield Vibrance Antifade Mounting Medium (Biozol, VEC-H-1700-10) and images were acquired with the Zeiss CellObserver with a Colibri LED illumination unit.

### Proximity-dependent biotinylation by APEX2

6 × 10^6^ cells were seeded per 15 cm dish 24 h prior to doxycycline induction of APEX2-fusion construct expression. 24 h after induction, culture medium was replaced and supplemented with 1 or 2.5 mM biotin-phenol (Iris Biotech, #LS-3500.1000) and incubated at 37 °C with 5% CO_2_ for 30 min. Following incubation, APEX2-labelling was initiated by addition of 30% H_2_O_2_ (CP26.1, Carl Roth) at a final concentration of 1 mM. The plate was gently agitated for exactly 1 min and the reaction was quenched by aspirating culture medium and washing three times with a quenching buffer (PBS with 10 mM sodium ascorbate, 10 mM sodium azide, 5 mM Trolox). For puromycin treatment, 1 mM puromycin (Invivogen, #ant-pr-1) was added in the BP-containing culture medium and was therefore incubated for 30 min prior to APEX2-labelling.

### Streptavidin enrichment of biotinylated proteins

After biotinylation, cells from 15 cm dishes were trypsinised, pelleted in quenching buffer, and washed once with PBS. Each pellet was resuspended in 100 μL 2× RIPA buffer and incubated on a rotating wheel for 90 min at 4 °C. Insoluble material was removed by centrifugation at 15,000xg for 30 min at 4 °C and the supernatant was transferred to a Zeba™ Spin Desalting Column, 7K MWCO (Thermo Fisher Scientific, #89882) to remove excess unreacted biotin-phenol. To enrich biotinylated proteins, 1 mg total cell lysate was incubated with 100 μL streptavidin magnetic sepharose beads (Cytiva, #28985799 ) equilibrated with RIPA buffer in a total volume of 1 mL. After 1 h incubation at RT on a rotating wheel, beads were washed twice with RIPA buffer, followed by 1 M KCl, 100 mM Na_2_CO_3_, 2 M urea in 10 mM Tris-HCl pH 8.0, and twice with RIPA buffer. Biotinylated proteins were eluted at 95 °C for 15 min in 3× SDS-PAGE sample buffer supplemented with 2 mM biotin and 20 mM DTT.

### Mass spectrometry data acquisition and label-free quantification

MTX2-APEX and APEX-TOMM22 samples were compared to their respective unbiotinylated control samples (H_2_O_2_ omitted) and to NES-APEX samples in three replicates. Additionally, MTX2-APEX and APEX-TOMM22 samples were analysed between puromycin-treated and -untreated conditions, in addition to APEX2-NES +/-puromycin samples in three replicates.

Proteins were purified by SDS-PAGE (4-12% NuPAGE tris gel (Invitrogen) for 7 min at 200 V and stained with Coomassie blue. The protein gel pieces were isolated and subjected to tryptic digestion as described previously ^28^.

Purified peptide samples were analysed on an Exploris 480 mass spectrometer (Thermo Fisher Scientific), connected directly to an Easy-nLC 1200 UHPLC or a Vanquish Neo UHPLC (Thermo Fisher Scientific). Peptides were separated on a 20-cm, 75–μm inner diameter analytical HPLC column (ID PicoTip fused silica emitter; New Objective, Berks, UK), packed in-house with ReproSil-Pur C18-AQ 1.9-μm silica beads (Dr Maisch GmbH, Ammerbuch, Germany). Separation was achieved with a 60-minute segmented gradient of solvents A (0.1% formic acid) and B (80% acetonitrile in 0.1% formic acid), at a flow rate of 200 nl/min. The column temperature was kept at 40°C using an integrated oven. Peptides were ionized through nanospray ionization with a capillary temperature of 275°C.

The mass spectrometer was operated in positive ion mode, and peptides were measured using data-dependent acquisition. Spectra were acquired over a scan range of 300 to 1750 m/z with an MS1 resolution of 60000. AGC was set to “custom” with a normalized value of 300% and an absolute target of 3.000e6. The top 20 most intense ions were selected for HCD fragmentation at 28% normalized collision energy, with a dynamic exclusion period of 30 seconds. Tandem MS (MS/MS) spectra were acquired at a resolution of 15,000.

### Mass-spectrometry data analysis

The raw data were processed with MaxQuant (version 2.2.0.0). The raw spectra were searched against the UniProt Homo sapiens database (104556 entries, downloaded on 30.01.2024) and common human contaminants. For MS and MS/MS, the peptide mass tolerances were set at 4.5 ppm and 20 ppm, respectively. Two missed cleavages were allowed for the tryptic digestion. Cysteine carbamidomethylation was used as a fixed modification, while N-terminal acetylation and methionine oxidation were set as variable modifications. The false discovery rate was maintained at 1% at both the peptide and protein levels. Both LFQ and Intensity-based absolute quantification (iBAQ) were enabled. All other parameters remained at default settings.

Downstream analysis of the ‘proteinGroups.txt’ output table was conducted in Perseus (version 2.0.10.0). Contaminants, reversed proteins, and proteins only identified by site were filtered out. Proteins were functionally annotated with Gene Ontology (GO) Biological Processes, GO Cellular Component, GO Molecular Functions, Kyoto Encyclopedia of Genes and Genomes, and MitoCarta3.0. Additionally, only proteins quantified in two out of three replicates in at least one group were kept in the dataset. iBAQ values were used and log10 transformed. Missing values were imputed separately for each column (width 0.3, down shift 1.8). A t-test was performed to identify significantly enriched proteins. Volcano plots of the -log10 p-values vs. the t-test differences were generated. Further visualization was performed using R (version 4.1.1) and GraphPad (version 8.0.1).

## Results

### APEX2 fusion proteins of TOMM22 and MTX2 are functional

TOM and SAM complexes must be in close vicinity since they cooperate to achieve integration of beta-barrel proteins into the mitochondrial outer membrane (MOM) ^6^. Nevertheless, the copy number of TOM complexes by far exceeds that of SAM complexes as demonstrated in yeast cells ^12, 13^, hence it can be expected that a fraction of TOM complexes differs in its set of proximal proteins from that of SAM. To map the proxisomes of the core TOM complex and SAM, we generated fusion proteins of TOMM22 and MTX2 and the proximity labelling enzyme APEX2 ^26^. To achieve a cytoplasmic location of the enzyme, APEX2 was fused to the N-terminus of TOMM22 or to the C-terminus of MTX2 (**Supporting Figure S1**; **Figure 1A**). We chose MTX2 over MTX1 or SAM50 because its C-terminus points towards the cytoplasm ^29^ and, in contrast to MTX1, it is not involved in a direct binding to the TOM complex ^11^, making it less likely that the addition of a large tag interferes with its function. In each case, both parts of the fusion proteins were separated by a ten amino acid long linker peptide and contained a V5 tag for detection of the protein ^22^.

**Fig. 1.**
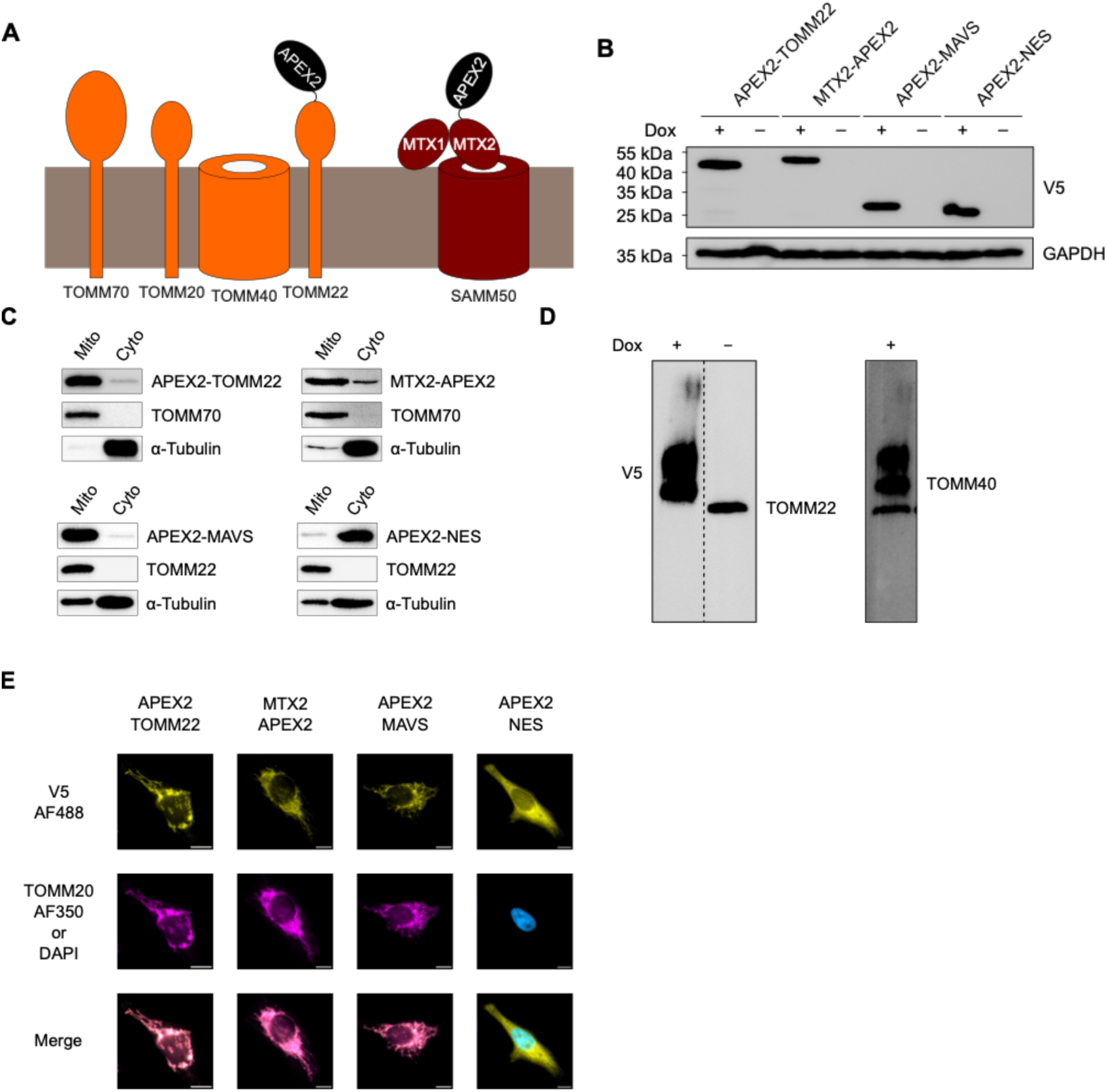
Tagging APEX2 on human TOMM22 and MTX2. (A) Schematic representation of TOMM22 and MTX2 tagged with APEX2 in their respective complexes at the MOM. (B) Western blot analysis of expression of each fusion construct upon 24 h doxycycline treatment. (C) Western blot analysis of subcellular localisation of each APEX2 fusion protein. Mitochondria and cytosol were isolated from each cell line via centrifugation after doxycycline treatment, TOMM70 or TOMM22 was used as a mitochondrial marker and α-tubulin was used as a cytosolic marker. (D) Incorporation of APEX2-TOMM22 into the TOM complex. Mitochondria from APEX2-TOMM22 with and without doxycycline treatment were isolated and solubilised before being subjected to BN-PAGE analysis, the left lane was stripped and re-probed with anti-TOMM40. (E) Fluorescence imaging of APEX2-TOMM22 -MTX2, - MAVS, and -NES fusion constructs. APEX2 fusion proteins were stained with anti-V5 antibody and Alexa Fluor 488 (yellow), endogenous TOMM20 was used as a mitochondrial marker and stained with Alexa Fluor 350 (magenta). For APEX2-NES, DAPI was used instead of a mitochondrial marker. Scale bar: 10 μm.

Expression cassettes that allow regulated expression under control of a doxycycline-dependent promoter were stably integrated into the genome of HeLa-EM2-11ht cells^30^. We also used the same master cell line to generate two control cell lines expressing a cytoplasmic APEX2 (APEX2-NES) and a MOM-targeted APEX2 (APEX2-MAVS). All four fusion proteins are expressed at similar levels upon 24 hr induction with doxycycline (**Figure 1B**). To minimise the effect of doxycycline on mitochondrial energetics and translation, we performed a test series with varying doxycycline concentrations to determine the lowest possible dosage for our purpose. At 0.3 μg/ul doxycycline, APEX2-TOMM22 was expressed at the same level as endogenous TOMM22, as detected by Western blot using an anti-TOMM22 antibody **(Supporting Figure S2A)**. At similar concentrations, we also observed sufficient expression of MTX2-APEX2 (**Supporting Figure S2B**). Expression of neither construct affected viability as indicated by a similar proliferation of cells expressing APEX2-TOMM22, MTX2-APEX2, or neither (**Supporting Figure S2C**).

To test if the APEX2 fusion proteins are correctly targeted, we performed subcellular fractionation and immunofluorescence staining (**Figure 1C and 1E**). APEX2 fusions of TOMM22 or MTX2, and the APEX2-MAVS control co-fractionated with a mitochondrial marker (TOMM70 or TOMM22; **Figure 1C**), whereas the majority of APEX2-NES co-fractionated with alpha-tubulin. Similar observations were made by indirect immunofluorescence, using an antibody against the V5 tag. In contrast to APEX2-NES, which showed a diffuse cytoplasmic staining, the other three proteins demonstrate a staining pattern overlapping with that of the mitochondrial marker TOMM20. These observations suggest a correct localization of APEX2-fused TOMM22 and MTX2. Further analysis by blue native gel electrophoresis (BN-PAGE) of the APEX2-TOMM22 fusion protein showed that it co-migrates with the TOMM40 pore, indicating correct integration into the TOM core complex (**Figure 1D**). To evaluate if the fusion proteins are functional, we performed knockdown experiments using siRNAs targeted only towards endogenous TOMM22 mRNAs. Western blot analysis demonstrated that knockdown of endogenous TOMM22 (**Supporting Figure S3A**) could be achieved to 84.6% compared to cells transfected with the negative control siRNA without affecting expression of the corresponding APEX2 fusion protein. Interestingly, following induction of MTX2-APEX2 expression, we observed a dramatic drop in the level of endogenous MTX2 (**Supporting Figure S3B**). Since the SAM complex is inherently a lot less abundant than the TOM complex, the expression level of MTX2 could be more tightly regulated either at the transcription level or by degradation of excess unassembled protein by the proteasome in order to maintain the precise stoichiometry of the multisubunit complex upon the increase in gene dosage ^31^.

Overall, cells lacking endogenous TOMM22 or MTX2 did not show any change in growth rate or morphology (**Supporting Figure S2C**) while the respective fusion protein was being expressed, indicating both APEX2 fusion constructs were able to functionally replace the endogenous ones.

### APEX2-TOMM22 and MTX2-APEX2 proxisomes overlap but differ

To define and compare the local protein environments of TOMM22 and MTX2 at the MOM, three independent replicates for proteins biotinylated via APEX2-TOMM22 and MTX2-APEX2 were analysed by bottom-up proteomics. During titration of various biotin-phenol concentrations we realized that similarly efficient biotinylation by APEX2-TOMM22 and MTX2-APEX2 is seen at 1mM, or 2.5 mM biotin-phenol, respectively (**Supporting Figure S4**). We therefore used these for the APEX2-TOMM22 replicates or for the MTX2-APEX2 replicates. In addition, we analysed six replicates for APEX2-NES, three at a concentration of 1 mM biotin-phenol and three at 2.5 mM biotin-phenol treatment. Furthermore, we included replicates of control experiments lacking H_2_O_2_. Biotinylation with BP and H₂O₂, quenching, lysis and capturing was essentially done according to previously published protocol (see Methods). The captured proteins were further measured by liquid chromatography-tandem mass spectrometry (LC-MS) after tryptic digestion. Missing values were imputed to increase the number of quantified proteins (see Methods). We identified up to 2,218 protein groups upon induction of the various fusion proteins and initiation of biotinylation by H₂O₂ (**Figure 2A**). Of these, up to 248 were annotated for mitochondrial localization by MitoCarta3.0 ^32^. Overall, we observed an excellent correlation between the replicates (**Suppl. Figure S5A**). We next evaluated the sub-organelle identifications of the captured biotinylated proteins (**Figure 2**), which showed that APEX2-TOMM22 and MTX2-APEX2 proxisomes had highly similar distributions, with comparable numbers of mitochondrial, MOM, MIM, IMS, and matrix proteins identified. Both constructs showed a clear enrichment of MOM proteins relative to the NES control. In contrast, matrix and MIM proteins were detected at similar levels across all three constructs and the unbiotinylated controls, suggesting a lack of matrix/MIM protein enrichment. Notably, MTX2-APEX2 recovered more RNA-binding proteins (71) than APEX2-TOMM22 (48), indicating potential differences in their local protein environments.

**Figure 2.**
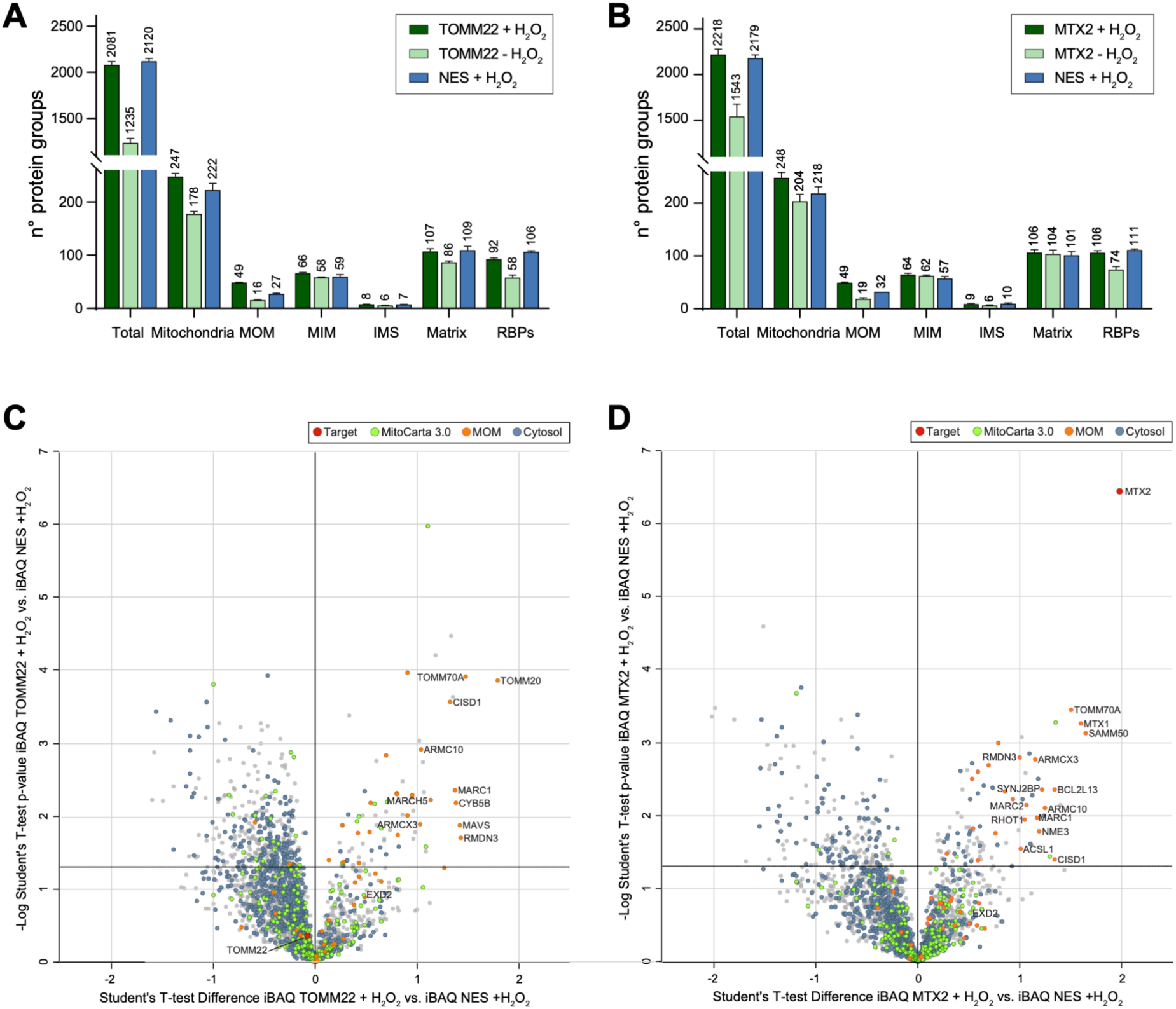
Proteomic analysis of APEX2-TOMM22 and MTX2-APEX2 proximity labeling. (A) Number of proteins identified in APEX2-TOMM22 +H₂O₂, APEX2-TOMM22 −H₂O₂, and APEX2-NES +H₂O₂ samples. (B) Number of proteins identified in MTX2-APEX2 +H₂O₂, MTX2-APEX2 −H₂O₂, and APEX2-NES +H₂O₂ samples. Proteins are categorized as total identified proteins, mitochondrial proteins, mitochondrial outer membrane (MOM), mitochondrial inner membrane (MIM), intermembrane space (IMS), matrix proteins, and RNA-binding proteins (RBPs). (C) Volcano plot of the APEX2-TOMM22 proxisome comparing APEX2-TOMM22 +H₂O₂ with APEX2-NES +H₂O₂ controls. The x-axis shows the Student’s t-test difference (log₂-transformed abundance ratio), and the y-axis shows the −log₁₀-transformed p-value. (D) Volcano plot of the MTX2-APEX2 proxisome comparing MTX2-APEX2 +H₂O₂ with APEX2-NES +H₂O₂ controls. The x-axis shows the Student’s t-test difference (log₂-transformed abundance ratio), and the y-axis shows the −log₁₀-transformed p-value. The target is highlighted in red. Mitochondrial proteins are shown in green, MOM proteins in orange, and cytosolic proteins in blue.

Besides the expected mitochondrial proteins, we also identified cytosolic proteins in the proxisomes of all three APEX2 fusion proteins. However, cytosolic proteins were depleted in the proxisomes of APEX2-TOMM22 and MTX2-APEX2 when directly compared to that of the cytosolic APEX2-NES (**Figure 2C and D**).

Of note, although MTX2 was the most enriched biotinylated protein in cells expressing MTX2-APEX2, this is not seen for TOMM22 in APEX2-TOMM22 cells. A potential explanation is the small number of aromatic amino acid residues that are generally targeted by the activated phenoxyl biotin radicals ^26^ in the cytoplasmic domain of TOMM22 (W49 and F55) ^33^, while the cytoplasmic part of MTX2 contains 25 aromatic amino acids ^29, 34^.

Preliminary analysis showed a specific enrichment of TOM complex proteins for APEX2-TOMM22 (TOMM20, TOMM70) and of SAM complex proteins for MTX2-APEX2 (MTX1, SAMM50), demonstrating that our proximal labelling was able to discriminate between distinct local environments despite the reported close spatial proximity of these protein complexes at the MOM ^10^.

### Core proteomic environments for TOMM22 and MTX2 remain largely unchanged upon translation inhibition

Due to increasing evidence for local translation in the mitochondrial vicinity, a major interest of our approach was to monitor the presence of RNA-binding proteins at different locations at the MOM. Given the established disruptive effect of puromycin on ongoing translation, we also asked whether puromycin treatment would alter the representation of RBPs as well as the overall proteomic architecture in the proxisome of TOMM22 and MTX2. Such changes have been observed before for subsets of mitochondrial-associated RBPs ^35^. However, we found that treatment with 200 µM puromycin for 30 min ^35^ prior to APEX2 labelling resulted in minimal alterations to the proxisomes of both APEX2-TOMM22 and MTX2-APEX2, suggesting only limited impact on the local protein environment under these conditions. To achieve more effective translation inhibition, the puromycin concentration was thus increased to 1 mM in subsequent experiments.

Comparison of the APEX2-TOMM22 proxisomes in the presence **(Figure 3, Supporting Figures S6 and S7)** and absence of puromycin revealed both a stable core set of proteins, as well as changes in the composition of the proximal proteome. A shared set of proteins, including TOMM70A, SYNJ2BP, MFN2, MFF, RHOT1, GDAP1, RMDN3, and CHCHD3, were detected under both conditions, indicating that TOMM22 remains stably associated with a specific MOM environment. In the absence of puromycin, the proximal proteome was more strongly enriched for MOM-, ER-, and MAM-associated proteins, including TOMM20, MAVS, MARCH5, ERP44, PDIA3, PDIA4, P4HB, ITPR3, and ERLIN1, consistent with a spatially restricted environment at mitochondrial import and membrane contact sites. This protein environment was lost upon puromycin treatment and replaced by an enrichment for proteins associated with inner mitochondrial compartments, respiratory chain function, and cytosolic regulatory pathways, including NDUFB10, CYC1, SLC25A46, BCL2L1, and VPS13A, indicating diminished spatial confinement and a broader detectable proteomic landscape.

**Figure 3.**
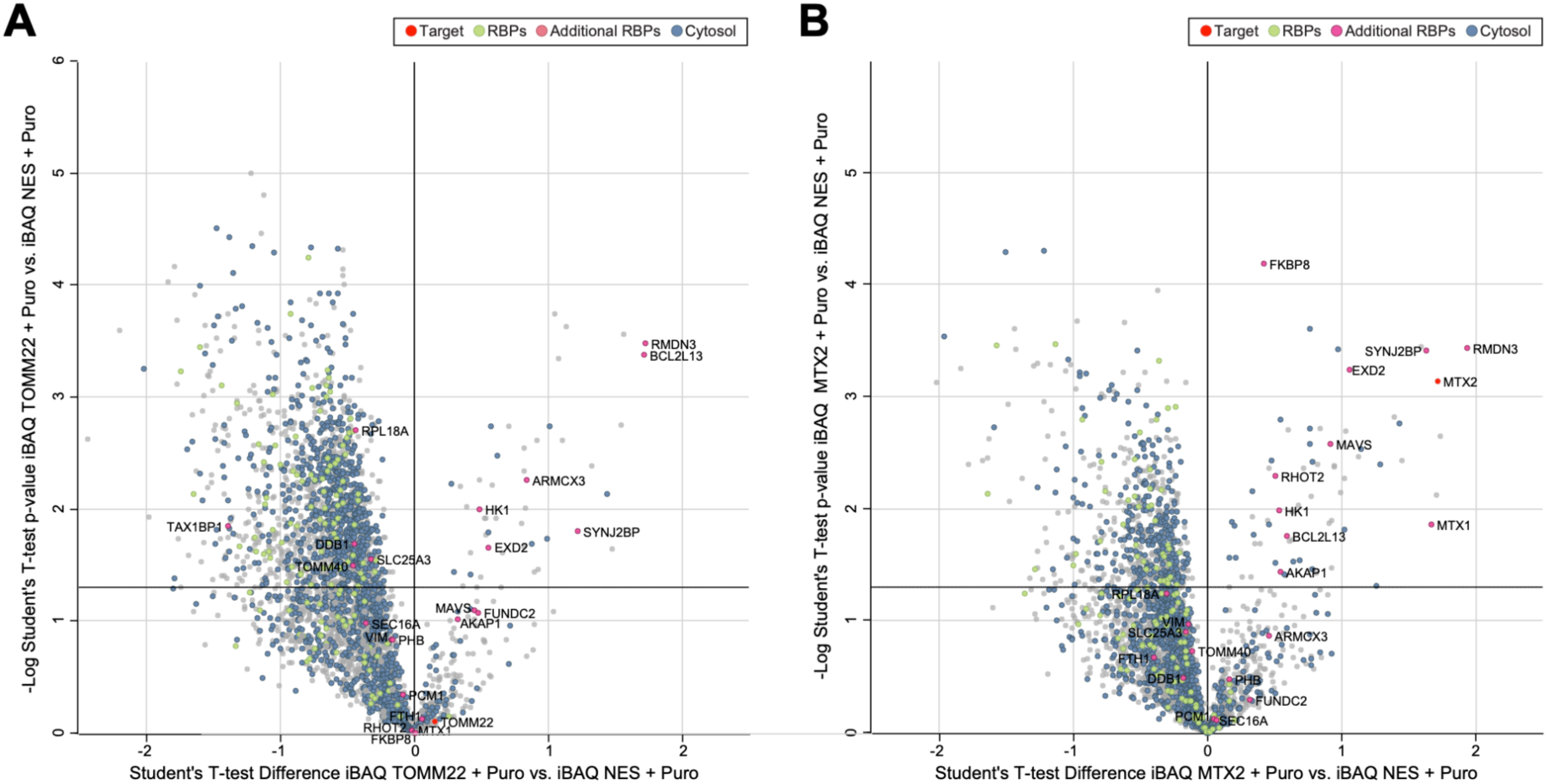
Analysis of TOMM22 and MTX2 proxisomes upon puromycin treatment. (A) Volcano plot of the APEX2-TOMM22 proxisome following puromycin treatment compared with the NES-APEX2 puromycin-treated control. X-axis shows the Student’s t-test difference (log₂-transformed iBAQ abundance ratio), y-axis shows the −log₁₀-transformed p-value. (B) Volcano plot of the MTX2-APEX2 proxisome following puromycin treatment compared with the NES-APEX2 puromycin-treated control. The x-axis shows the Student’s t-test difference (log₂-transformed iBAQ abundance ratio), and the y-axis shows the −log₁₀-transformed p-value. The target is highlighted in red. RBPs are shown in green, manually selected additional RBPs in orange, and cytosolic proteins in blue

A similar pattern was observed for MTX2, although with a more diverse proteomic composition. Puromycin treatment led to a pronounced redistribution of proteins within the MTX2-associated proximal proteome while preserving a stable core mitochondrial environment. A substantial subset of proteins, including the components of the β-barrel assembly machinery (MTX1, MTX2, SAMM50), TOMM70A, as well as factors linked to mitochondrial architecture (IMMT, CHCHD3), dynamics (MFN1, MFF, RHOT2), and specific RNA-associated processes (SYNJ2BP, EXD2), were consistently detected under both conditions. This defines a stable, translation-independent MTX2-associated environment that spans multiple layers of mitochondrial organization. In contrast, the untreated condition was characterized by a broader and more heterogeneous set of proteins, including numerous regulatory and RNA-associated factors such as ZFP36L2, ZC3H11A (both RNA-associated proteins), ZCCHC11, ELP6 (both regulatory proteins that modify RNA), and SRPK1 (an SR-RBP specific kinase), indicating that under steady-state conditions MTX2 samples a complex and functionally diverse cellular interface. Following puromycin treatment, additional proteins were detected that were functionally linked to mitochondrial respiration, including COX4I1, CYCS, and CYC1, as well as proteins involved in mitochondrial membrane organization or dynamics, such as SLC25A46, VPS13A, and MYO19.

### EXD2 is a novel TOM/SAM associated RNA binding protein

Several RBPs have recently been reported to localize in proximity to, or associate with, the MOM or TOM complex ^18, 35–37^. Among these, SYNJBP2 was identified in proximity-labelling studies using both MAVS-APEX2 and TOMM20-APEX2 constructs ^22, 35^. Another well-characterized MOM-associated RBP is AKAP1, which has been identified in multiple studies as an RNA-binding scaffold protein at the mitochondrial surface ^18, 20, 36^. More recently, AKAP1 was shown to bind mRNAs encoding components of the electron transport chain independently of active translation, thereby promoting localized translation and facilitating mitochondrial protein import ^18^. Together, these findings indicate that TOM-associated regions of the MOM may participate in the spatial organization of mRNA localization and localized translation at the mitochondrial surface.

To investigate whether RBPs were similarly enriched in our TOMM22-APEX2 and MTX2-APEX2 datasets, proteins were annotated using the RBPDB database ^38^. Compared to the APEX2-NES control, two RBPs (AKAP1 and IREB2) were enriched in the APEX2-TOMM22 dataset, while five RBPs (AKAP1, ANKHD1, ZC3HAV1L, ZFP36L2, and ZC3H11A) were enriched in the MTX2-APEX2 dataset. However, we noted that the relatively stringent RBPDB annotation appeared incomplete, as several proteins more recently characterized as RBPs, including SYNJ2BP ^39^, were absent from the database. Following additional manual annotation of highly enriched candidates, four proteins were selected for further validation: EXD2 ^40^, AKAP1 ^20^, RMDN3 and ARMCX3 ^35^.

To determine whether these candidates physically associate with TOM/SAM-related microenvironments, co-immunoprecipitation was performed using V5-tagged APEX2-TOMM22 and MTX2-APEX2 constructs. RMDN3, AKAP1, and ARMCX3 did not co-immunoprecipitate with either TOMM22-APEX2 or MTX2-APEX2 **(Figure 4A and 4B, Supporting Figure S8)**, suggesting that these proteins may either interact only weakly or too transiently with TOM and SAM or reside more broadly within the surrounding proteomic landscape without direct association. Notably, despite its established role as a MOM-associated RBP and its enrichment in our mass spectrometric datasets, AKAP1 did not show detectable physical interaction with TOMM22 or MTX2 under the tested conditions. In contrast, EXD2 co-immunoprecipitated with both APEX2-TOMM22 and MTX2-APEX2, indicating a substantially stronger and potentially more specific association.

**Figure 4.**
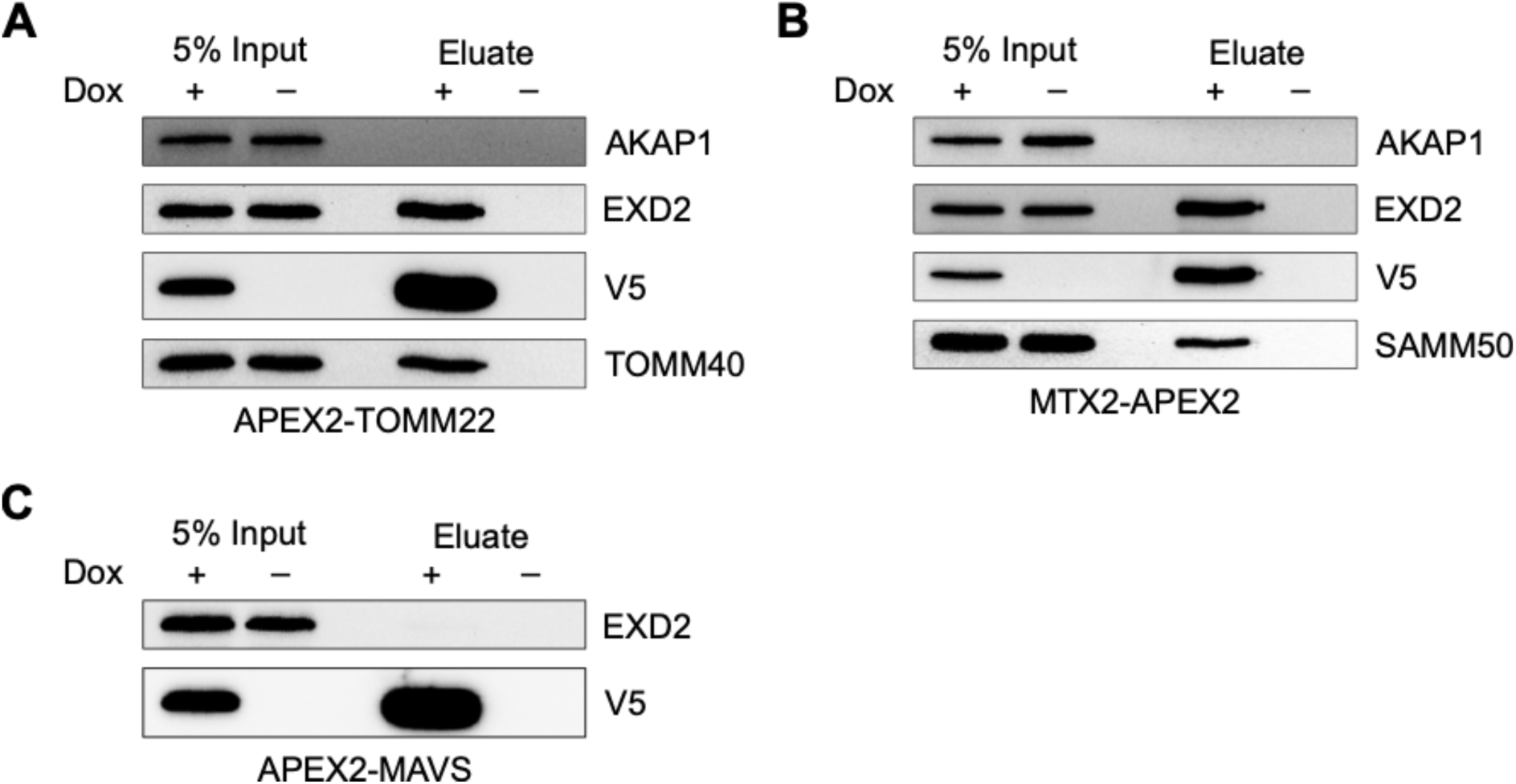
**Co-immunoprecipitation of EXD2 with APEX2-TOMM22 and MTX2-APEX2**. (A) and (B) Western blot analysis of anti-V5 co-immunoprecipitation with solubilized mitochondria from APEX2-TOMM22 and MTX2-APEX2 cell lines. TOMM40 or SAMM50 was used as their respective positive control and AKAP1 was probed as negative control for both. (C) Mitochondria were isolated and solubilized from the APEX2-MAVS cell line and subjected to anti-V5 co-immunoprecipitation to check for interaction with EXD2.

To assess whether this interaction merely reflected the general presence of EXD2 at the mitochondrial outer membrane, we additionally tested a V5-tagged MOM-localized APEX2-MAVS construct ^41^. In contrast to TOMM22-APEX2 and MTX2-APEX2, no co-IP signal was detected with APEX2-MAVS **(Figure 4C)**. These findings suggest that the observed interaction is not mediated by the V5 epitope or the APEX2 enzyme itself but is instead dependent on the interaction with TOMM22- or MTX2-associated environment at the MOM. Furthermore, EXD2 enrichment was consistently observed across biological replicates and remained detectable following puromycin treatment, indicating that its association with TOM/SAM-associated regions is maintained independently of ongoing translation.

## Discussion

Proximity labelling by the engineered ascorbate peroxidase APEX2 has emerged as a powerful approach for the study of suborganellar proteomes and transcriptomes. The fast-labelling kinetics of APEX2, combined with the membrane impermeability and short half-life of the biotin-phenoxyl radical, allow highly specific, spatially restricted biotinylation and therefore precise interrogation of local molecular environments. In this study, fusion of APEX2 to the cytosolic domains of TOMM22 and MTX2 enabled comprehensive mapping of the proximal proteomes associated with two major MOM protein complexes, TOM and SAM.

Importantly, as the SAM complex is considerably less abundant than the TOM complex and is involved in the biogenesis of only a subset of mitochondrial proteins, MTX2-APEX2 required substantially higher BP concentrations to achieve efficient labelling. Consistent with this, the MTX2-APEX2 proxisome appeared more diverse and contained a larger proportion of cytosolic proteins. This might be due to a more pronounced escape of biotin-phenoxyl radicals to react with cytosolic proteins. Alternatively, it may reflect the lower local protein density and more spatially dispersed molecular environment surrounding MTX2, resulting in less efficient but still spatially specific labelling of SAM-associated regions at the MOM.

Beyond the identification of established components of the import machineries including TOMM20 and TOMM70 for APEX2-TOMM22 and MTX1 and SAMM50 for MTX2-APEX2, several RBPs were detected in proximity to both TOM and SAM complexes. These findings are in line with previous proximity labelling studies at the MOM ^18, 35^, supporting the emerging concept that the MOM serves as a platform for localized RNA translation. Among the tested RBPs, we confirmed the association of EXD2 with both TOM and SAM, whereas AKAP1, despite its established role in local translation of mitochondrial proteins ^18^, did not display comparable enrichment. Moreover, the persistence of EXD2 in puromycin-treated samples suggests that its association with mitochondrial import receptors is largely translation-independent.

Although EXD2 was previously proposed to localize to the nucleus and mitochondrial matrix, more recent studies have identified it at the MOM ^40, 42^. Furthermore, Sandoz et al. ^43^ provided functional insight into this MOM-localized pool of EXD2 by demonstrating that, upon UV-induced genotoxic transcriptional stress, EXD2 translocates from the MOM to the nucleus, where it promotes RNA polymerase II backtracking through degradation of nascent mRNA, thereby facilitating transcription restart after DNA repair.

On the other hand, while previous studies identified EXD2 at the MOM, its precise localization within the MOM and its potential interacting partners remained unclear. Our findings provide further support for the MOM localization of EXD2 and additionally suggest that EXD2 may not be uniformly distributed across the mitochondrial surface but rather enriched within specific TOM- and SAM-associated microdomains. In co-immunoprecipitation, EXD2 was identified to be associated with APEX2-TOMM22 and MTX2-APEX2, but not with APEX2-MAVS, arguing against a generalized distribution across the MOM. This is in line with findings by Yoo and Rhee ^42^, in which EXD2 was also detected in a TOMM20-APEX2 proximity-labelling dataset. However, the function of EXD2 at the MOM and the molecular mechanism governing its stress-dependent relocalisation remain important questions for future investigation. Together, our findings provide insight into the spatial organization of mitochondrial protein import sites and further support the association of RBPs with TOM- and SAM-associated regions at the MOM, which may provide local sites for the enrichment of factors involved in broader cellular regulatory processes.

## Acknowledgement

We thank the team of the Proteome Center Tübingen for sample processing and for technical support in data processing. We also thank Birgit Singer-Krüger (IFIB) for project discussions.

## Data Availability

The mass spectrometry proteomics data will be available on PRIDE as soon as the upload is complete.

## Transparency and Ethics Reporting

This research did not involve human or animal participants.

## Author Contributions

XW and RPJ wrote the paper with the support of CCR and BM. XW and RPJ designed the project. XW performed all experiments. CCR, with support by BM, was responsible for MS sample processing, data curation and analysis, and visualization. Figures were made by XW and CCR. RPJ and BM provided resources, supervision, and funding.

## Supporting Information

**Supporting Figure S1.**
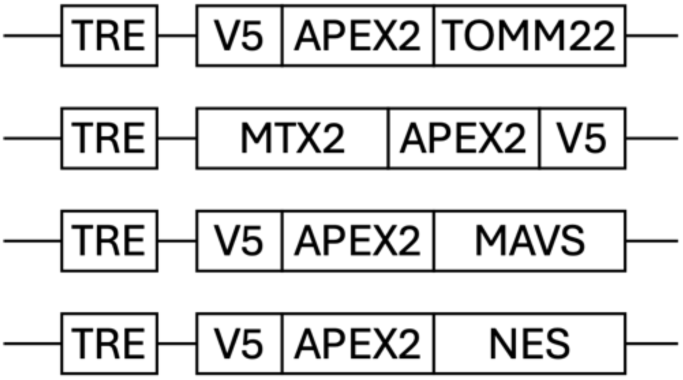
**Expression constructs used in this study**. TRE: tetracyclin-responsive element with bidirectional minimal CMV promoter; V5: V5 peptide epitope tag; MAVS: mitochondrial outer membrane targeting peptide from Mitochondrial Antiviral-Signalling protein; NES: nuclear export signal.

**Supporting Fig. S2.**
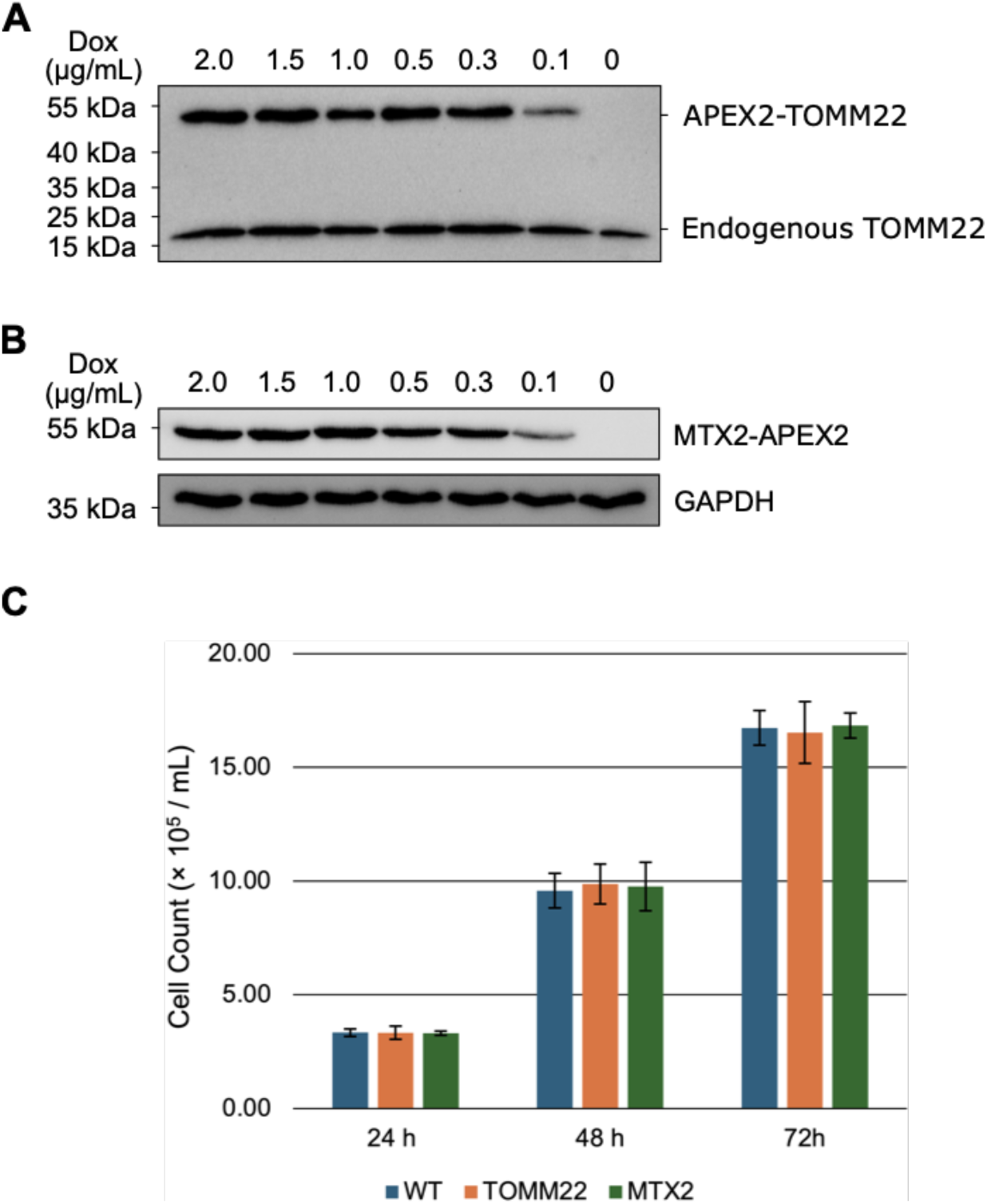
Expression of APEX2-TOMM22 and MTX2-APEX2. **(A)** and **(B)** Various doxycycline concentrations were tested to induce expression of APEX2-TOMM22 and -MTX2. **(C)** Cell growth analysis of wild-type (WT), APEX2-TOMM22 and MTX2-APEX2 cell lines. 1× 10^6^ cells were seeded per 100 mm dish per cell line and were treated with doxycycline for 24 to 72 h, before trypsinisation, resuspension in 6 mL culture medium, and counting. Data represent mean ± SD from *n* = 3 independent experiments.

**Supporting Fig. S3.**
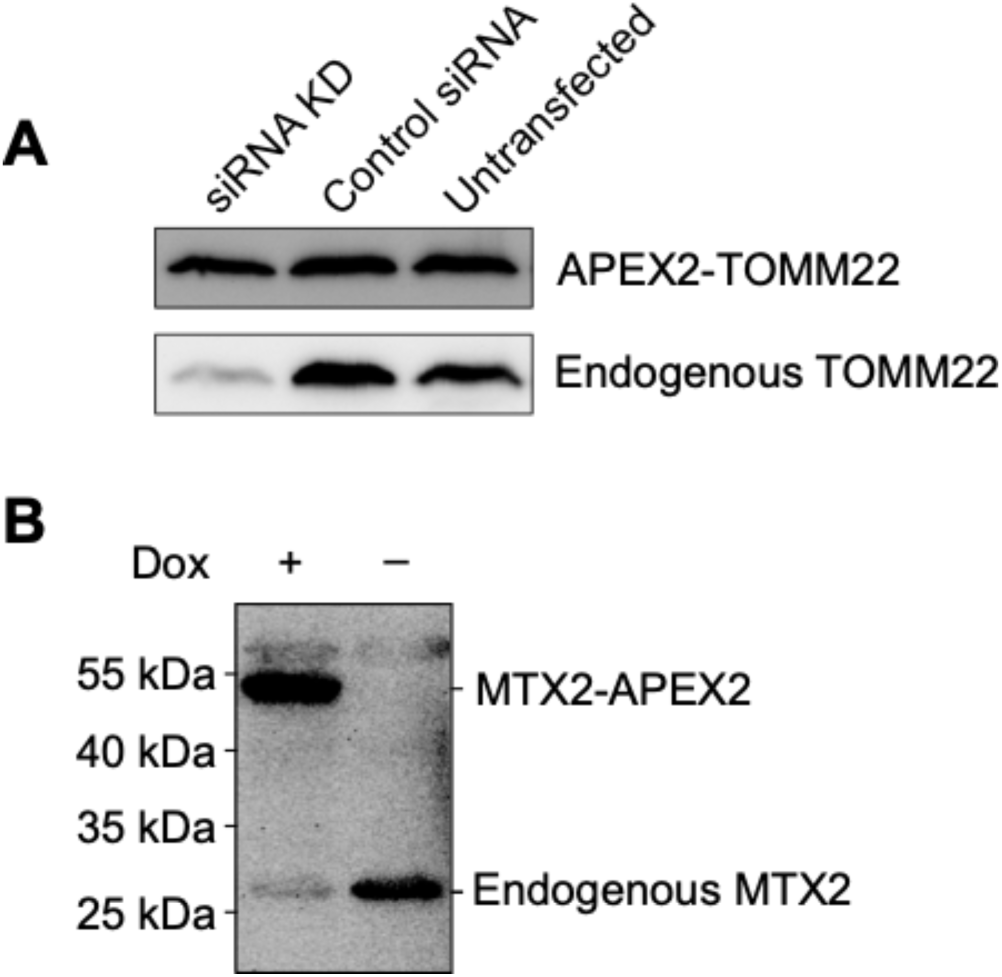
Functional replacement of TOMM22 and MTX2 by their APEX2-tagged variants. **(A)** Cells expressing APEX2-TOMM22 were treated with siRNA targeting endogenous TOMM22 and protein depletion was assessed by immunoblotting. Despite efficient reduction of the endogenous TOMM22 signal, cells showed no detectable impairment in cell growth or viability, indicating that the APEX2-TOMM22 fusion protein could functionally replace the endogenous protein. **(B)** Upon expression of MTX2-APEX2, expression level of endogenous MTX2 was markedly reduced as assessed by immunoblotting. Despite the reduction, MTX2-APEX2-expressing cells showed no detectable defect in cell growth, suggesting that MTX2-APEX2 could functionally compensate for reduced endogenous MTX2 levels.

**Supporting Fig. S4.**
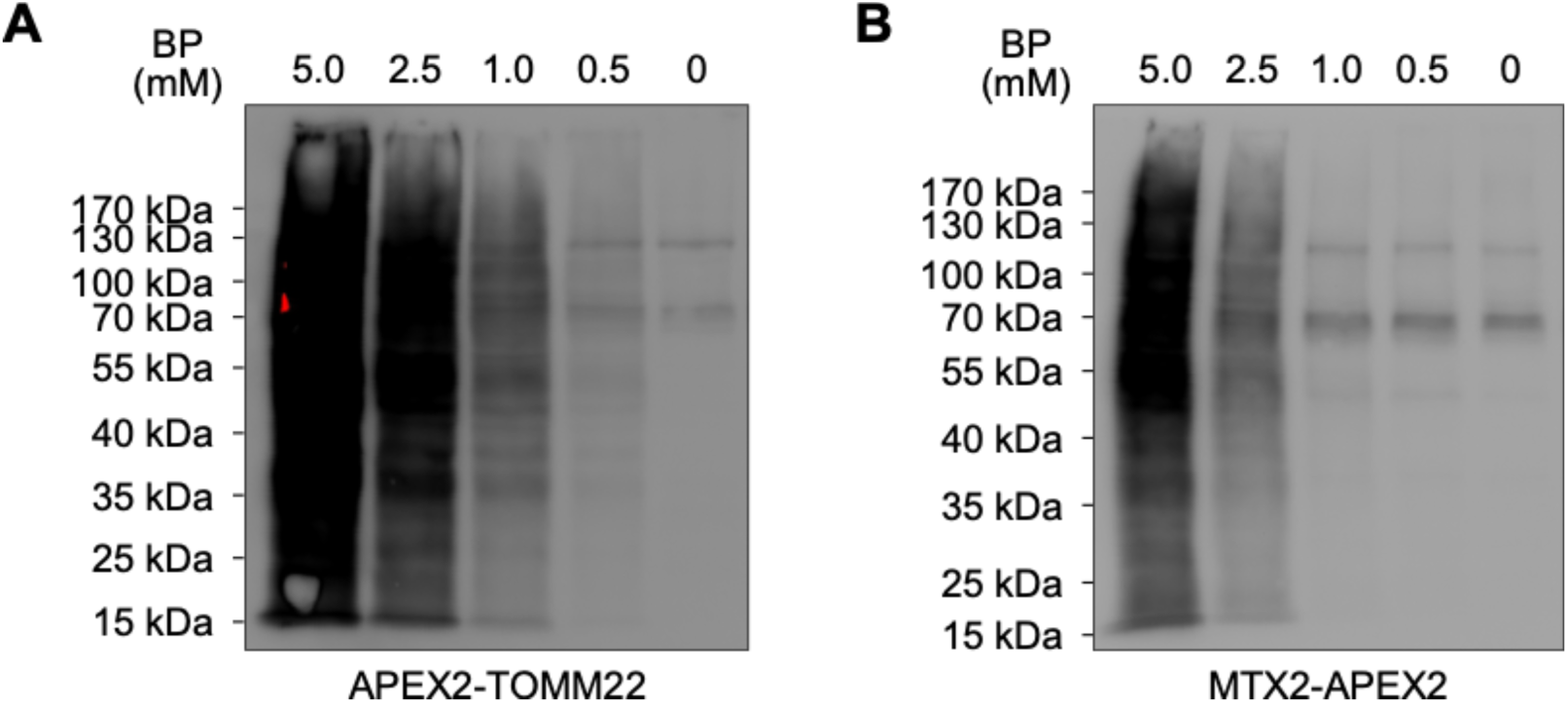
Biotin-phenol titration for APEX2-TOMM22 and MTX2-APEX2. **(A)** and **(B)** Various biotin-phenol concentrations were tested for optimal biotinylation signal in each cell line. Biotinylated proteins were detected by streptavidin-HRP.

**Supporting Fig. S5.**
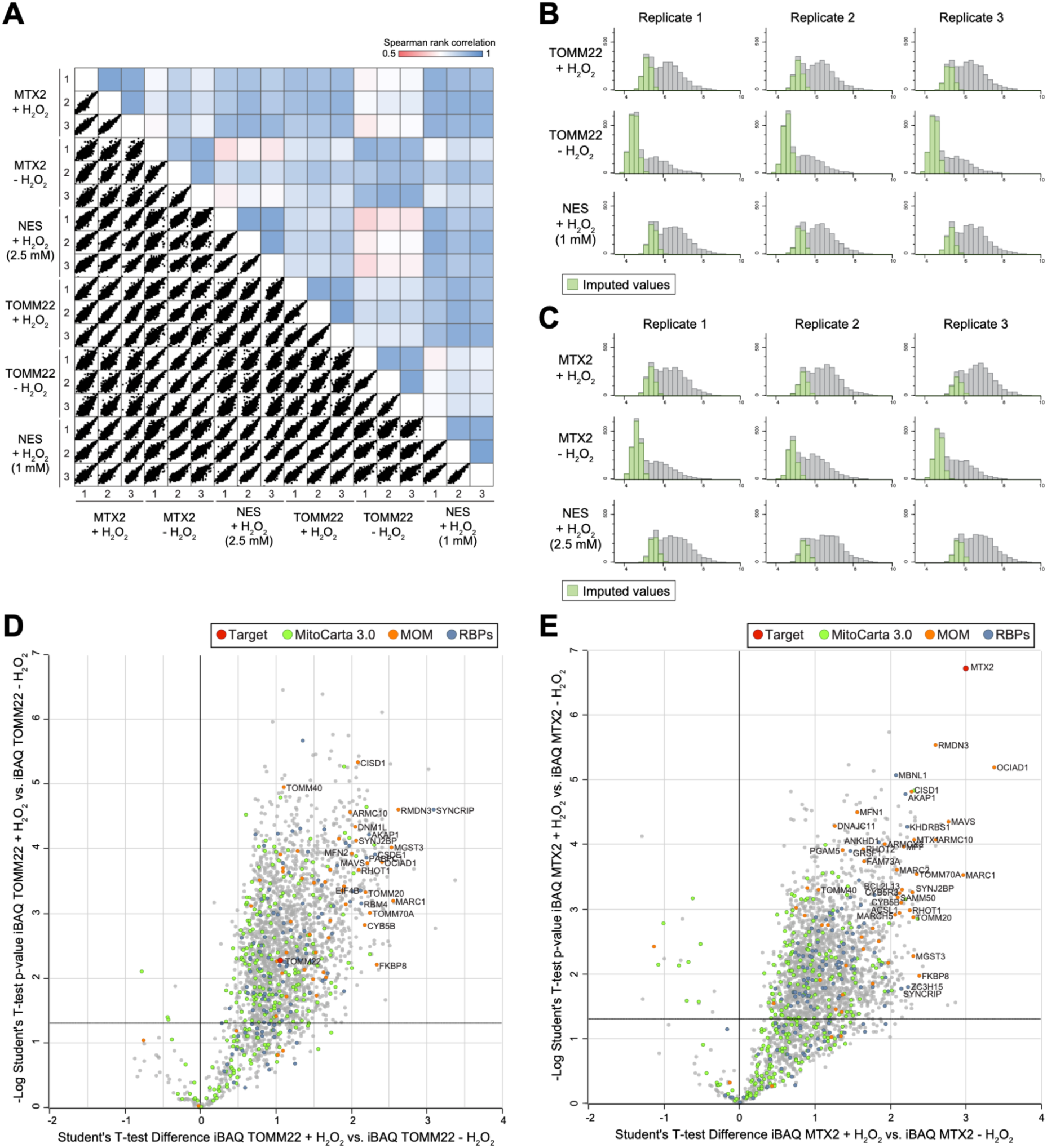
Analysis of the proxisomes in the APEX2-TOMM22 and MTX2-APEX2 datasets. **(A)** Spearman rank correlation comparing all biological replicates and experimental conditions. (**B)** Distribution plots showing imputation of missing values for APEX2-TOMM22 datasets across three biological replicates. (**C)** Distribution plots showing imputation of missing values for MTX2-APEX2 datasets across three biological replicates. Imputed values (green) represent predominantly low-abundance proteins replaced using a normal distribution-based imputation method. (**D)** Volcano plot of the APEX2-TOMM22 proxisome comparing APEX2-TOMM22 +H₂O₂ with APEX2-TOMM22 −H₂O₂ controls. X-axis shows the Student’s t-test difference (log₂-transformed iBAQ abundance ratio), and the Y-axis shows the −log₁₀-transformed p-value. (**E)** Volcano plot of the MTX2-APEX2 proxisome comparing MTX2-APEX2 +H₂O₂ with MTX2-APEX2 −H₂O₂ controls. X-axis shows the Student’s t-test difference (log₂-transformed iBAQ abundance ratio), and Y-axis shows the −log₁₀-transformed p-value. The target is highlighted in red. Mitochondrial proteins are shown in green, MOM proteins in orange, and RBPs in blue.

**Supporting Fig. S6.**
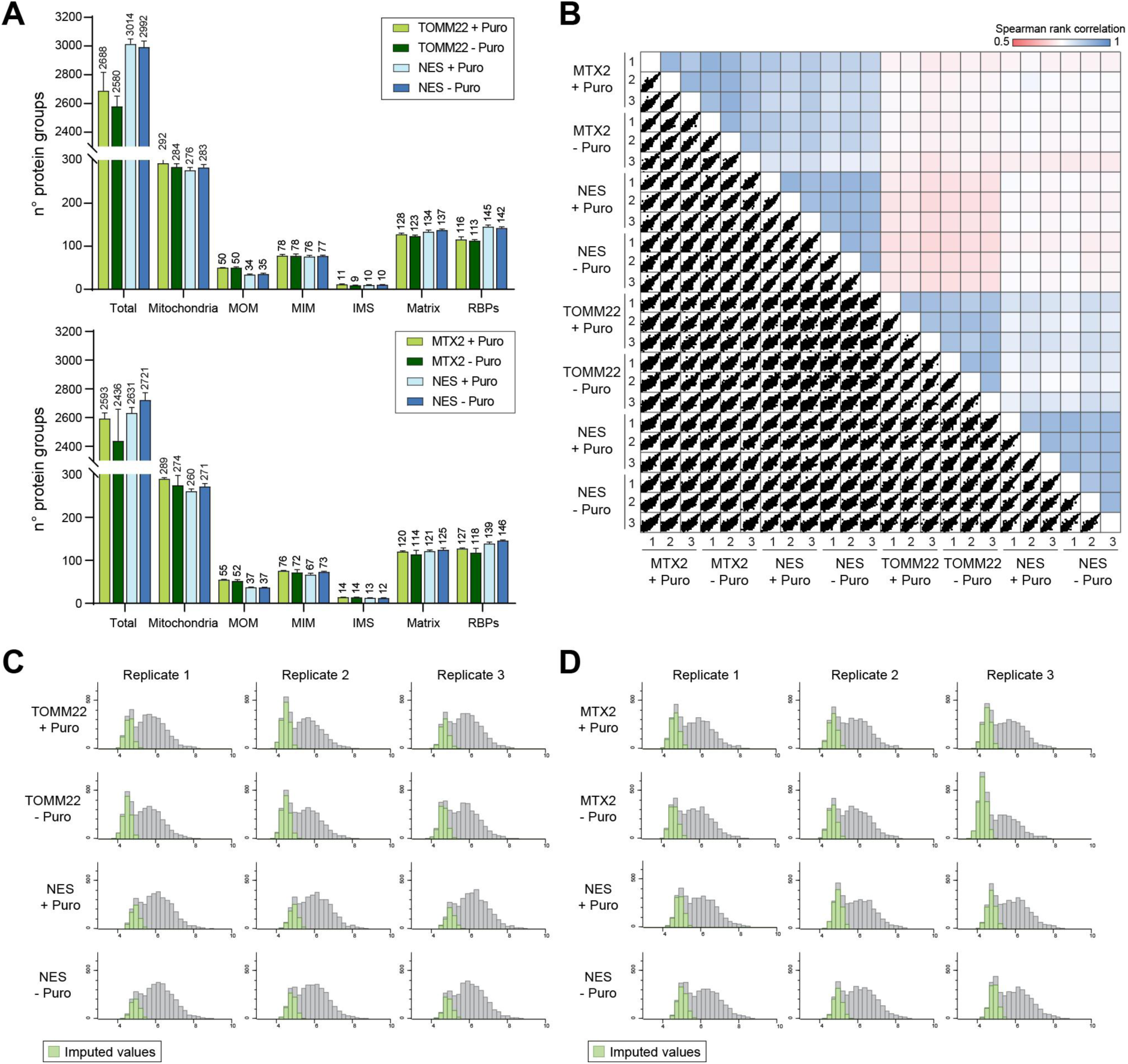
Proteomic quality control of the proxisomes of TOMM22 and MTX2 analysis after translational inhibition by puromycin. (A) In the upper graph, number of proteins identified in APEX2-TOMM22 + Puromycin, APEX2-TOMM22 − Puromycin, APEX2-NES + Puromycin and APEX2-NES - Puromycin samples. In the upper graph, number of proteins identified in MTX2-APEX2 + Puromycin, MTX2-APEX2 − Puromycin, APEX2-NES + Puromycin and APEX2-NES - Puromycin samples. Proteins are categorized as total identified proteins, mitochondrial proteins, mitochondrial outer membrane (MOM), mitochondrial inner membrane (MIM), intermembrane space (IMS), matrix proteins, and RNA-binding proteins (RBPs). (B) Spearman rank correlation comparing all biological replicates and experimental conditions. (C) Distribution plots showing imputation of missing values for APEX2-TOMM22 datasets across three biological replicates. (D) Distribution plots showing imputation of missing values for MTX2-APEX2 datasets across three biological replicates. Imputed values (green) represent predominantly low-abundance proteins replaced using a normal distribution-based imputation method.

**Supporting Fig. S7.**
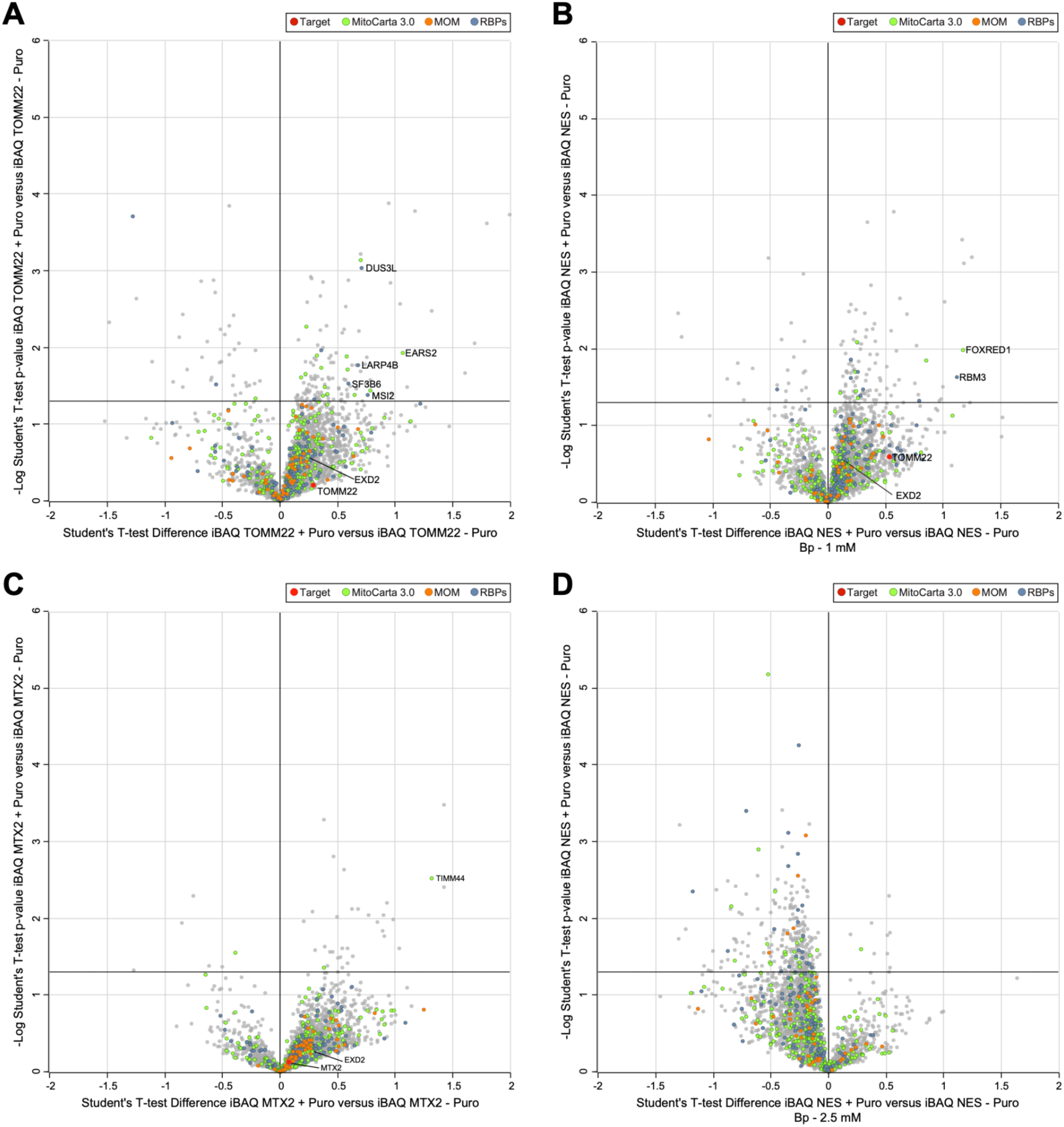
Proteomic analysis of the proxisomes of TOMM22 and MTX2 after translational inhibition by puromycin.(A) Volcano plot comparing APEX2-TOMM22 samples treated with puromycin (+Puro) and untreated controls (−Puro). (**B)** Volcano plot comparing APEX2-NEScontrol samples treated with puromycin (+Puro) and untreated controls (−Puro) with 1 µM BP **C.** Volcano plot comparing MTX2-APEX2 samples treated with puromycin (+Puro) and untreated controls (−Puro). (**D)** Volcano plot comparing NES-APEX2 control samples treated with puromycin (+Puro) and untreated controls (−Puro) with 2.5 µM BP. The X-axis shows the Student’s *t*-test difference (log₂-transformed iBAQ abundance ratio), and the Y-axis shows the −log₁₀-transformed *p*-value. The target is highlighted in red. Mitochondrial proteins are shown in green, MOM proteins in orange, and cytosolic proteins in blue.

**Supporting Figure S8.**
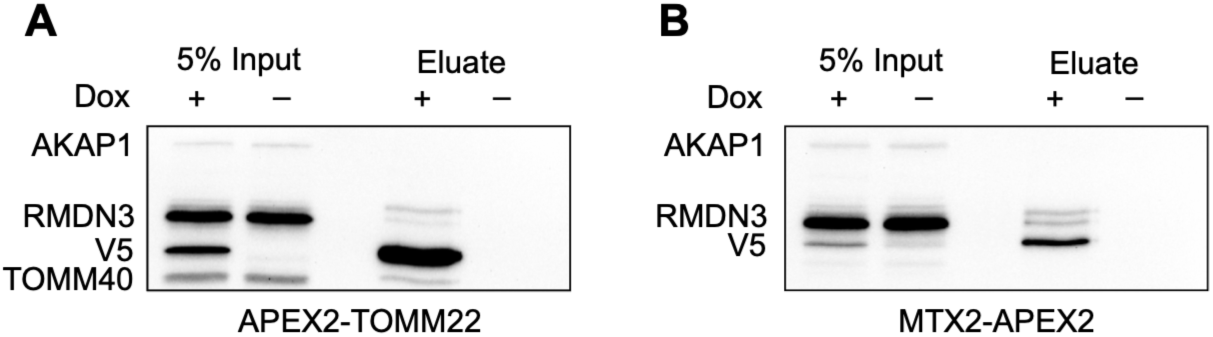
**Lack of co-immunoprecipitation of RMDN3 with APEX2-TOMM22 and MTX2-APEX2**. (A) and (B) Western blot analysis of anti-V5 co-immunoprecipitation with solubilized mitochondria from APEX2-TOMM22 and MTX2-APEX2 cell lines. TOMM40 was used as their respective positive control and AKAP1 was probed as negative control for both.

**Supporting Table 1:**
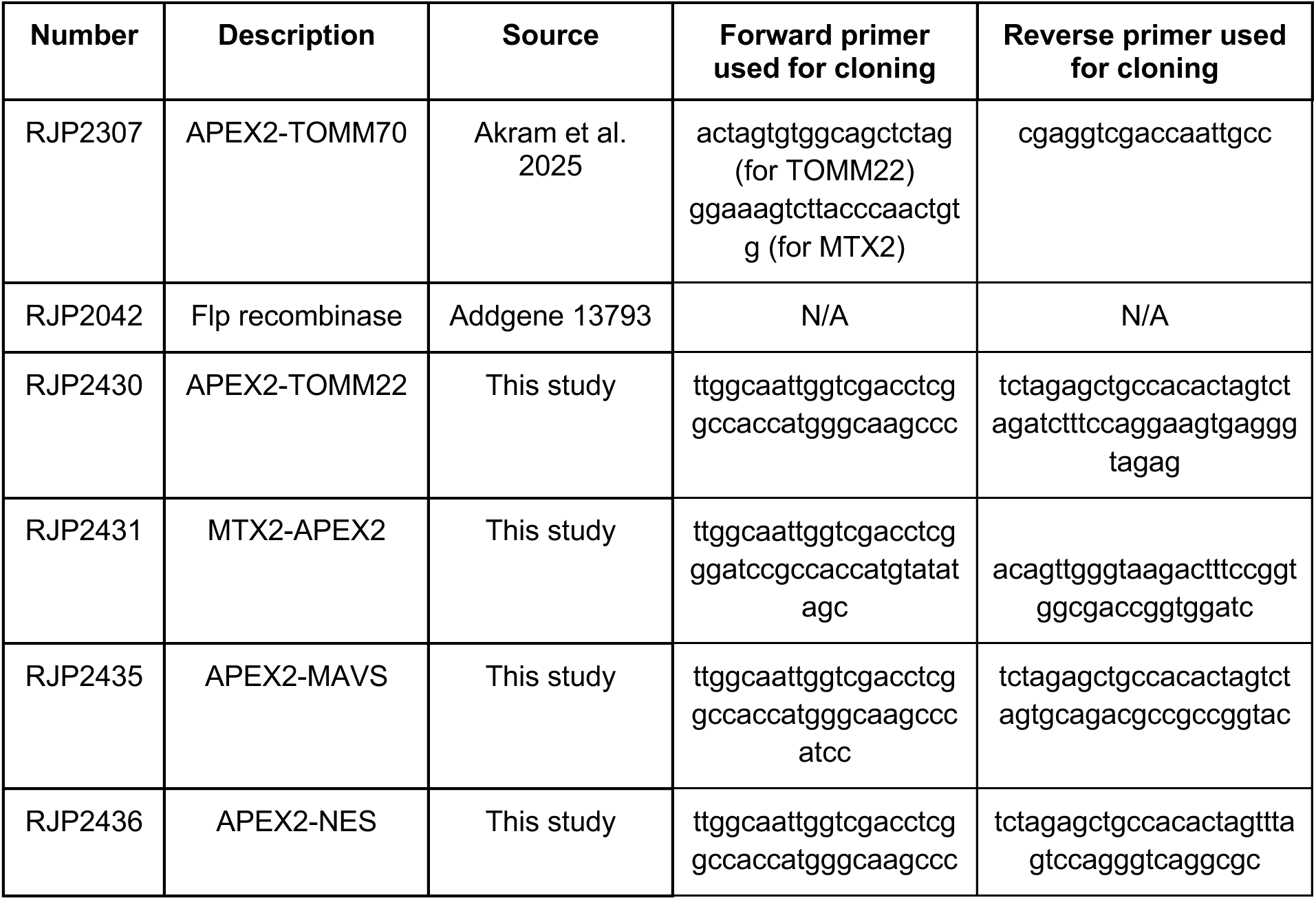
Plasmids used in this study.

**Supporting Table 2:** Antibodies used in this study.

| **Antibody** | **Company** | **Catalogue Number** | **Use** | **Dilution** |
| --- | --- | --- | --- | --- |
| mouse anti-V5 | Thermo Fisher | R960-25 | WB, IF | 1:5,000 (WB)<br>1:1,000 (IF) |
| rabbit anti-V5 | Cell Signaling Technology | 13202 | WB | 1:1,000 |
| rabbit anti-TOMM20 | Proteintech | 11802-1-AP | IF | 1:200 |
| rabbit anti-TOMM22 | Abcam | ab246862 | WB | 1:5,000 |
| rabbit anti-TOMM40 | Abcam | ab185543 | WB | 1:2,000 |
| mouse anti-TOMM70 | Proteintech | 14528-1-AP | WB | 1:500 |
| rabbit anti-MTX2 | Abcam | ab272607 | WB | 1:500 |
| rabbit anti-SAM50 | Abcam | ab167430 | WB | 1:5,000 |
| rabbit anti-AKAP1 | Proteintech | 15618-1-AP | WB | 1:2,000 |
| rabbit anti-EXD2 | Proteintech | 20138-1-AP | WB | 1:500 |
| mouse anti-GAPDH | Proteintech | 60004-1-Ig | WB | 1:20,000 |
| mouse anti-α-tubulin | Sigma-Aldrich | T6199 | WB | 1:500 |
| rabbit anti-ARMCX3 | Proteintech | 25705-1-AP | WB | 1:1,000 |
| rabbit anti-RMDN3 | Proteintech | 20641-1-AP | WB | 1:1,000 |
| streptavidin-HRP | Invitrogen | S911 | WB | 0.5 μg/mL |
| sheep anti-mouse HRP | Jackson ImmunoResearch | 515-035-003 | WB | 1:10,000 |
| goat anti-rabbit HRP | Millipore | AP307P | WB | 1:10,000 |
| goat anti-mouse Alexa Fluor 488 | Invitrogen | A11001 | IF | 1:200 |
| donkey anti-rabbit Alexa Fluor 350 | Invitrogen | A10039 | IF | 1:100 |

## References

(1) Suomalainen, A.; Nunnari, J. Mitochondria at the crossroads of health and disease. Cell 2024, 187 (11), 2601–2627. DOI: 10.1016/j.cell.2024.04.037.

(2) Baker, Z. N.; Forny, P.; Pagliarini, D. J. Mitochondrial proteome research: the road ahead. Nat Rev Mol Cell Biol 2024, 25 (1), 65–82. DOI: 10.1038/s41580-023-00650-7 From NLM Medline.

(3) Wiedemann, N.; Pfanner, N. Mitochondrial Machineries for Protein Import and Assembly. Annu Rev Biochem 2017, 86, 685–714. DOI: 10.1146/annurev-biochem-060815-014352.

(4) Becker, T.; Song, J.; Pfanner, N. Versatility of Preprotein Transfer from the Cytosol to Mitochondria. Trends Cell Biol 2019, 29 (7), 534–548. DOI: 10.1016/j.tcb.2019.03.007.

(5) Araiso, Y.; Endo, T. Structural overview of the translocase of the mitochondrial outer membrane complex. Biophys Physicobiol 2022, 19, e190022. DOI: 10.2142/biophysico.bppb-v19.0022 From NLM PubMed-not-MEDLINE.

(6) Diederichs, K. A.; Pitt, A. S.; Varughese, J. T.; Hackel, T. N.; Buchanan, S. K.; Shaw, P. L. Mechanistic insights into fungal mitochondrial outer membrane protein biogenesis. Curr Opin Struct Biol 2022, 74, 102383. DOI: 10.1016/j.sbi.2022.102383 From NLM Medline.

(7) Hohr, A. I.; Straub, S. P.; Warscheid, B.; Becker, T.; Wiedemann, N. Assembly of beta-barrel proteins in the mitochondrial outer membrane. Biochim Biophys Acta 2015, 1853 (1), 74–88. DOI: 10.1016/j.bbamcr.2014.10.006 From NLM Medline.

(8) Endo, T.; Yamano, K. Transport of proteins across or into the mitochondrial outer membrane. Biochim Biophys Acta 2010, 1803 (6), 706–714. DOI: 10.1016/j.bbamcr.2009.11.007 From NLM Medline.

(9) Kozjak-Pavlovic, V.; Ross, K.; Benlasfer, N.; Kimmig, S.; Karlas, A.; Rudel, T. Conserved roles of Sam50 and metaxins in VDAC biogenesis. EMBO Rep 2007, 8 (6), 576–582. DOI: 10.1038/sj.embor.7400982 From NLM Medline.

(10) Qiu, J.; Wenz, L. S.; Zerbes, R. M.; Oeljeklaus, S.; Bohnert, M.; Stroud, D. A.; Wirth, C.; Ellenrieder, L.; Thornton, N.; Kutik, S.;, et al. Coupling of mitochondrial import and export translocases by receptor-mediated supercomplex formation. Cell 2013, 154 (3), 596–608. DOI: 10.1016/j.cell.2013.06.033 From NLM Medline.

(11) Wenz, L. S.; Ellenrieder, L.; Qiu, J.; Bohnert, M.; Zufall, N.; van der Laan, M.; Pfanner, N.; Wiedemann, N.; Becker, T. Sam37 is crucial for formation of the mitochondrial TOM-SAM supercomplex, thereby promoting beta-barrel biogenesis. J Cell Biol 2015, 210 (7), 1047–1054. DOI: 10.1083/jcb.201504119 From NLM Medline.

(12) Pfanner, N.; Warscheid, B.; Wiedemann, N. Mitochondrial proteins: from biogenesis to functional networks. Nat Rev Mol Cell Biol 2019, 20 (5), 267–284. DOI: 10.1038/s41580-018-0092-0 From NLM Medline.

(13) Morgenstern, M.; Stiller, S. B.; Lubbert, P.; Peikert, C. D.; Dannenmaier, S.; Drepper, F.; Weill, U.; Hoss, P.; Feuerstein, R.; Gebert, M.;, et al. Definition of a High-Confidence Mitochondrial Proteome at Quantitative Scale. Cell Rep 2017, 19 (13), 2836–2852. DOI: 10.1016/j.celrep.2017.06.014.

(14) Cohen, B.; Golani-Armon, A.; Arava, Y. S. Emerging implications for ribosomes in proximity to mitochondria. Semin Cell Dev Biol 2024, 154 (Pt B), 123–130. DOI: 10.1016/j.semcdb.2023.01.003 From NLM Publisher.

(15) Craig, E. A. Hsp70 at the membrane: driving protein translocation. BMC Biol 2018, 16 (1), 11. DOI: 10.1186/s12915-017-0474-3 From NLM Medline.

(16) Zilio, E.; Schlegel, T.; Zaninello, M.; Rugarli, E. I. The role of mitochondrial mRNA translation in cellular communication. J Cell Sci 2025, 138 (9). DOI: 10.1242/jcs.263753 From NLM Medline.

(17) Zhu, Z.; Mallik, S.; Stevens, T. A.; Huang, R.; Levy, E. D.; Shan, S. O. Principles of cotranslational mitochondrial protein import. Cell 2025, 188 (20), 5605–5617 e5614. DOI: 10.1016/j.cell.2025.07.021 From NLM Medline.

(18) Luo, J.; Khandwala, S.; Hu, J.; Lee, S. Y.; Hickey, K. L.; Levine, Z. G.; Harper, J. W.; Ting, A. Y.; Weissman, J. S. Proximity-specific ribosome profiling reveals the logic of localized mitochondrial translation. Cell 2025. DOI: 10.1016/j.cell.2025.08.002 From NLM Publisher.

(19) Garcia-Rodriguez, L. J.; Gay, A. C.; Pon, L. A. Puf3p, a Pumilio family RNA binding protein, localizes to mitochondria and regulates mitochondrial biogenesis and motility in budding yeast. J Cell Biol 2007, 176 (2), 197–207. DOI: 10.1083/jcb.200606054 From NLM Medline.

(20) Gabrovsek, L.; Collins, K. B.; Aggarwal, S.; Saunders, L. M.; Lau, H. T.; Suh, D.; Sancak, Y.; Trapnell, C.; Ong, S. E.; Smith, F. D.;, et al. A-kinase-anchoring protein 1 (dAKAP1)-based signaling complexes coordinate local protein synthesis at the mitochondrial surface. J Biol Chem 2020, 295 (31), 10749–10765. DOI: 10.1074/jbc.RA120.013454.

(21) Harbauer, A. B.; Hees, J. T.; Wanderoy, S.; Segura, I.; Gibbs, W.; Cheng, Y.; Ordonez, M.; Cai, Z.; Cartoni, R.; Ashrafi, G.;, et al. Neuronal mitochondria transport Pink1 mRNA via synaptojanin 2 to support local mitophagy. Neuron 2022, 110 (9), 1516–1531 e1519. DOI: 10.1016/j.neuron.2022.01.035 From NLM Medline.

(22) Akram, S.; Zittlau, K. I.; Sharma, K.; Fitzgerald, J. C.; Rafiq, N.; Macek, B.; Jansen, R. P. Proximity Labeling Reveals RNA-Binding Proteins Associating with the Human Mitochondrial Import Receptor TOMM20. J Proteome Res 2026, 25 (2), 1055–1070. DOI: 10.1021/acs.jproteome.5c00905 From NLM Medline.

(23) Meurant, S.; Mauclet, L.; Dieu, M.; Arnould, T.; Eyckerman, S.; Renard, P. Endogenous TOM20 Proximity Labeling: A Swiss-Knife for the Study of Mitochondrial Proteins in Human Cells. Int J Mol Sci 2023, 24 (11). DOI: 10.3390/ijms24119604 From NLM Medline.

(24) Qin, W.; Cho, K. F.; Cavanagh, P. E.; Ting, A. Y. Deciphering molecular interactions by proximity labeling. Nat Methods 2021, 18 (2), 133–143. DOI: 10.1038/s41592-020-01010-5.

(25) Samavarchi-Tehrani, P.; Samson, R.; Gingras, A. C. Proximity Dependent Biotinylation: Key Enzymes and Adaptation to Proteomics Approaches. Mol Cell Proteomics 2020, 19 (5), 757–773. DOI: 10.1074/mcp.R120.001941.

(26) Lam, S. S.; Martell, J. D.; Kamer, K. J.; Deerinck, T. J.; Ellisman, M. H.; Mootha, V. K.; Ting, A. Y. Directed evolution of APEX2 for electron microscopy and proximity labeling. Nat Methods 2015, 12 (1), 51–54. DOI: 10.1038/nmeth.3179 From NLM Medline.

(27) Schagger, H.; von Jagow, G. Blue native electrophoresis for isolation of membrane protein complexes in enzymatically active form. Anal Biochem 1991, 199 (2), 223–231. DOI: 10.1016/0003-2697(91)90094-a From NLM Medline.

(28) Shevchenko, A.; Tomas, H.; Havlis, J.; Olsen, J. V.; Mann, M. In-gel digestion for mass spectrometric characterization of proteins and proteomes. Nat Protoc 2006, 1 (6), 2856–2860. DOI: 10.1038/nprot.2006.468 From NLM Medline.

(29) Diederichs, K. A.; Ni, X.; Rollauer, S. E.; Botos, I.; Tan, X.; King, M. S.; Kunji, E. R. S.; Jiang, J.; Buchanan, S. K. Structural insight into mitochondrial beta-barrel outer membrane protein biogenesis. Nat Commun 2020, 11 (1), 3290. DOI: 10.1038/s41467-020-17144-1 From NLM Medline.

(30) Weidenfeld, I.; Gossen, M.; Low, R.; Kentner, D.; Berger, S.; Gorlich, D.; Bartsch, D.; Bujard, H.; Schonig, K. Inducible expression of coding and inhibitory RNAs from retargetable genomic loci. Nucleic Acids Res 2009, 37 (7), e50. DOI: 10.1093/nar/gkp108 From NLM Medline.

(31) Harper, J. W.; Bennett, E. J. Proteome complexity and the forces that drive proteome imbalance. Nature 2016, 537 (7620), 328–338. DOI: 10.1038/nature19947 From NLM Medline.

(32) Rath, S.; Sharma, R.; Gupta, R.; Ast, T.; Chan, C.; Durham, T. J.; Goodman, R. P.; Grabarek, Z.; Haas, M. E.; Hung, W. H. W.;, et al. MitoCarta3.0: an updated mitochondrial proteome now with sub-organelle localization and pathway annotations. Nucleic Acids Res 2021, 49 (D1), D1541–D1547. DOI: 10.1093/nar/gkaa1011 From NLM Medline.

(33) Su, J.; Liu, D.; Yang, F.; Zuo, M. Q.; Li, C.; Dong, M. Q.; Sun, S.; Sui, S. F. Structural basis of Tom20 and Tom22 cytosolic domains as the human TOM complex receptors. Proc Natl Acad Sci U S A 2022, 119 (26), e2200158119. DOI: 10.1073/pnas.2200158119 From NLM Medline.

(34) Armstrong, L. C.; Saenz, A. J.; Bornstein, P. Metaxin 1 interacts with metaxin 2, a novel related protein associated with the mammalian mitochondrial outer membrane. J Cell Biochem 1999, 74 (1), 11–22. From NLM.

(35) Qin, W.; Myers, S. A.; Carey, D. K.; Carr, S. A.; Ting, A. Y. Spatiotemporally-resolved mapping of RNA binding proteins via functional proximity labeling reveals a mitochondrial mRNA anchor promoting stress recovery. Nat Commun 2021, 12 (1), 4980. DOI: 10.1038/s41467-021-25259-2.

(36) Sharma, S.; Fazal, F. M. Localization of RNAs to the mitochondria-mechanisms and functions. RNA 2024, 30 (6), 597–608. DOI: 10.1261/rna.079999.124 From NLM Medline.

(37) Vardi-Oknin, D.; Arava, Y. Characterization of Factors Involved in Localized Translation Near Mitochondria by Ribosome-Proximity Labeling. Frontiers in Cell and Developmental Biology 2019, 7, 305. DOI: 10.3389/fcell.2019.00305 From NLM PubMed-not-MEDLINE.

(38) Cook, K. B.; Kazan, H.; Zuberi, K.; Morris, Q.; Hughes, T. R. RBPDB: a database of RNA-binding specificities. Nucleic Acids Res 2011, 39 (Database issue), D301–308. DOI: 10.1093/nar/gkq1069 From NLM Medline.

(39) Mullari, M.; Lyon, D.; Jensen, L. J.; Nielsen, M. L. Specifying RNA-Binding Regions in Proteins by Peptide Cross-Linking and Affinity Purification. J Proteome Res 2017, 16 (8), 2762–2772. DOI: 10.1021/acs.jproteome.7b00042 From NLM Medline.

(40) Hensen, F.; Moretton, A.; van Esveld, S.; Farge, G.; Spelbrink, J. N. The mitochondrial outer-membrane location of the EXD2 exonuclease contradicts its direct role in nuclear DNA repair. Sci Rep 2018, 8 (1), 5368. DOI: 10.1038/s41598-018-23690-y From NLM Medline.

(41) Fazal, F. M.; Han, S.; Parker, K. R.; Kaewsapsak, P.; Xu, J.; Boettiger, A.; Chang, H. Y.; Ting, A. Y. APEX-seq: RNA subcellular localization by proximity labeling. 2019. DOI: 10.21203/rs.2.1857/v1.

(42) Yoo, C. M.; Rhee, H. W. APEX, a Master Key To Resolve Membrane Topology in Live Cells. Biochemistry 2020, 59 (3), 250–259. DOI: 10.1021/acs.biochem.9b00785 From NLM Medline.

(43) Sandoz, J.; Cigrang, M.; Zachayus, A.; Catez, P.; Donnio, L. M.; Elly, C.; Nieminuszczy, J.; Berico, P.; Braun, C.; Alekseev, S.;, et al. Active mRNA degradation by EXD2 nuclease elicits recovery of transcription after genotoxic stress. Nat Commun 2023, 14 (1), 341. DOI: 10.1038/s41467-023-35922-5 From NLM Medline.

